# SAMPLE: an R package to estimate sampling effort for species’ occurrence rates

**DOI:** 10.1101/2024.06.10.598212

**Authors:** Henrique Bravo, Yacine Ben Chehida, Sancia E.T. van der Meij

## Abstract

Species’ occurrence rates are the backbone of many ecological studies. Sampling of species occurrence, however, can come with challenges and might prove more difficult than anticipated. Logistical difficulties, limited funds or time, elusiveness or rarity of species and difficult sampling environments are all examples of scenarios that might contribute to (undesired) small sample sizes. In order to help circumvent some of these difficulties and uncertainties, we present `SAMPLE`, an `R` package that aims to inform the user whether the amount of sampling conducted is enough to accurately estimate the occurrence rate of species. We use a simulation approach to help verify the accuracy of the package and to help guide the user in choosing the most appropriate values for the available parameters. Moreover, we provide a real data set where we used `SAMPLE` to estimate the occurrence rate of various coral-dwelling species on their hosts and the minimum number of samples required for an accurate estimation. This provided example data set includes closely related host species, single or multiple symbionts on a single host coral taxon, and data points obtained from different depths to illustrate how occurrence rates can vary depending on the provided input. Due to its simplicity and easiness of use, this package allows for users to run it whilst in the field to estimate if sampling is sufficient or if the sampling approach needs to be adapted for a particular species. We hope that this package proves itself useful to users that need to estimate occurrence or prevalence rates of species and do not always have the possibility to obtain large sample sizes.

## Introduction

Sample size is an important factor in any ecological research project. Large sample sizes improve the precision of estimations and the power of statistical tests, but also increases the costs and length of a field campaign (Underwood, 1997; Singh and Masuku, 2014). Study design typically includes estimations of sampling effort based on experience (Kenkel et al., 1989; Schreiber and Brauns, 2010), experimental design (Bernstein and Zalinski, 1983; Underwood and Chapman, 2003) and possibly, data simulations (Nadon and Stirling, 2006; Guerra-Castro et al., 2021).

To study biodiversity patterns, one of the most frequent types of ecological data collected is the occurrence of species (i.e. presence/absence) and how it relates to environmental and geographical patterns (Soberón and Peterson, 2004). This data can be useful, for example, towards studies on: habitat suitability of species (Hirzel et al., 2006), biogeography (Hanski, 1982), conservation (Rondinini et al., 2006), or community ecology (Gotelli, 2004). Prevalence (rate) is a measure of occurrence (commonly used in parasitology and disease ecology), and used here as the mean proportion of host taxa inhabited by a symbiont (Bush et al., 1997).

A large sample size is usually preferable when it comes to accurately determining the occurrence or prevalence rate of species (Gregory and Blackburn, 1991; Gotelli and Colwell, 2001). The relationship between sample size and accurate estimation of rates is not, however, a linear one and it might not be necessary to carry out an exhaustive sampling effort to obtain a sample size that is representative of the studied community (Gregory and Woolhouse, 1993; Chao and Bunge, 2002). The uncertainty of occurrence and prevalence rates of species rapidly decreases after a minimum number of samples has been reached (e.g. 10-20 individuals) and stabilises shortly after (Underwood, 1997; Jovani and Tella, 2006). In situations where it can be difficult to increase the sampling effort due to, for example, limited time or funds, logistical difficulties or elusiveness of study species, it is essential to know whether the effort carried out is enough or not. There are statistical methods that have been developed to cope with reduced or unbalanced sample sizes (Tella et al., 1999; Paterson and Lello, 2003; Jovani and Tella, 2006). Moreover, specific statistical approaches and `R` packages have been developed to predict the amount of sampling effort needed in a study, but they tend to require the provision of input data that will help in that sampling effort’s estimation (Anderson and Santana-Garcon, 2015; Guerra-Castro et al., 2021).

Given the usefulness, but also the difficulty, of knowing what the necessary sampling effort is for having accurate occurrence/prevalence rates of species without providing any initial input, we developed an `R` package to help with such estimations: `SAMPLE`. This package allows the user to determine whether the amount of sampling conducted so far is sufficient, or whether specific taxa are underrepresented in the data set. It has the added value that due to its simplicity and ease of use, it can be run whilst in the field. All of the parameters used in this package have set default values that should be the most useful for the majority of users, but they can be manually adjusted to fit the different needs of users and their study environments.

This package can be used for all kinds of terrestrial and marine community assemblages; however, as an example, we simulated data on hosts/symbionts to estimate the prevalence rates of the symbionts on their hosts in order to verify the accuracy in prevalence rate estimation of the `SAMPLE R` package. This simulation takes known prevalence rates of symbionts in populations of different sizes and tries to estimate the prevalence rates of these populations using different combinations of parameters (e.g. replicate number, different number of successive points).

Additionally, this paper uses a real data set of different species of obligate coral-dwelling symbionts of stony corals and hydrocorals to highlight if the prevalence rate of the symbionts across different coral host species was accurately estimated, and if so, what would have been the minimum sampling needed.

### Functionality of SAMPLE

The `SAMPLE R` package presented here takes as input a data set that can be accessed directly in the package or in Supplementary material (S3), where each row corresponds to a single data point. In the example provided prevalence rates are calculated with the names of species in the first column corresponding to the hosts, and its symbiont(s)’ occurrence(s) noted down with 1 (i.e. presence) and 0 (i.e. absence) in the remainder of the columns. To estimate, for example, the occurrence rates of species across habitats, the first column should contain the species of choice, and the remainder of the columns should denote the different habitats with 1 (i.e. presence) and 0 (i.e. absence) as values.

The data set is subsequently exposed to a process of random sampling with replacement, where a sample of *k* individuals from the initial data set is randomly sampled and its prevalence rate calculated with *k* ranging between 1 and *k*_*MAX*_ (i.e. the maximum number of available individuals). The process is repeated *n* times (user defined, default *n* = 50) in order to obtain an average occurrence or prevalence rate with *k* samples over *n* iterations.

The stability point of the occurrence or prevalence rate (i.e. the minimum number of samples needed to accurately determine the occurrence or prevalence rate of the study system) is determined by first computing the difference of *x* successive (mean) prevalence rates (user defined, default = 10) that are below a threshold *y* (user defined, default *y* = 2). Because *y* values become smaller as *k* increases, *y* is divided by the square root of *k* (i.e. 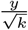) to correct for the effect of sample size. The difference between the minimum and maximum values among the *x* means (Delta: Δ) is then computed. The stability point is set as the first of the *x* successive means below a *z* threshold (user defined, default *z* = 1).

A schematic representation of this explanation can be found in Fig. 1. This is then repeated for each host and each symbiont species, with the plot being generated at the end (see section on Visualisation and Fig. 2 for an example).

**Fig. 1.**
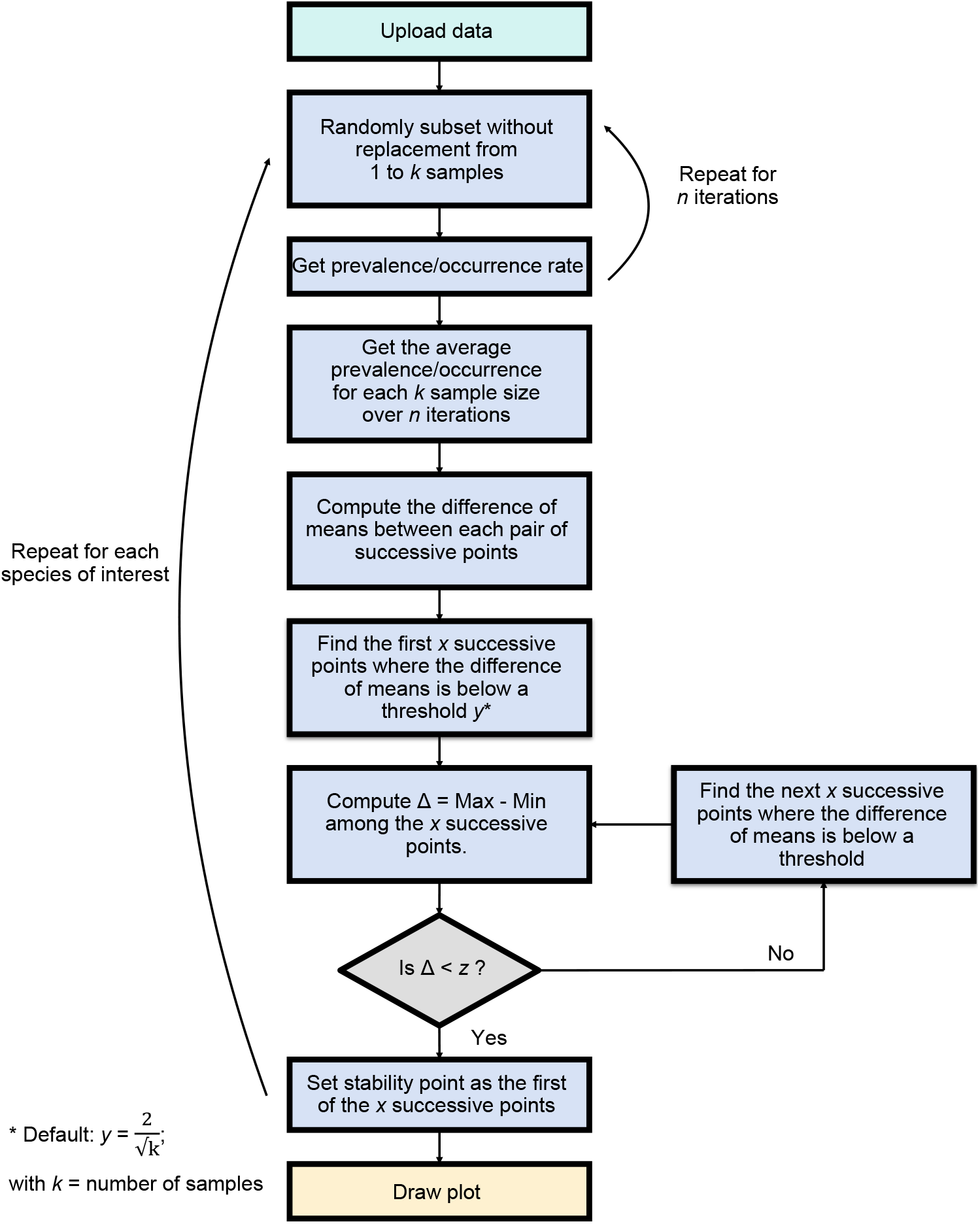
Schematic representation of the functionality of the `SAMPLE R` package.

**Fig. 2.**
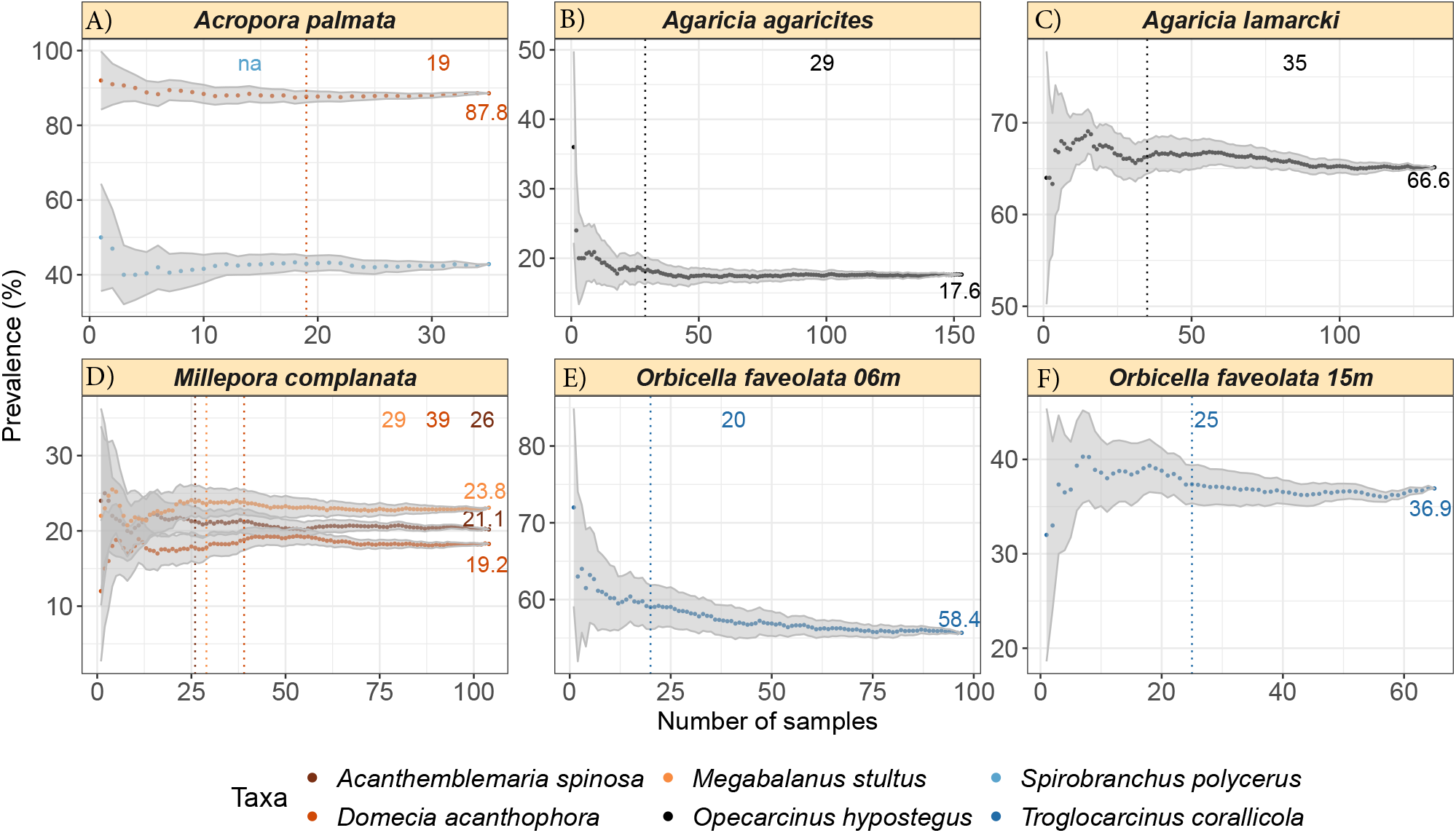
The minimum number of samples needed for the estimation of stable prevalence rates of different species of symbionts on their respective coral hosts. For a more detailed explanation of the different elements on the plots, please refer to Fig. 3.

### Simulated data: prevalence rates in a host/symbiont system

We simulated a host/symbiont data set with known population sizes and prevalence rates in order to test the validity and accuracy of the `SAMPLE R` package. Three population sizes were chosen (i.e. 100, 1000 and 10000) and prevalence rates of 10-90% with increments of 10% were then calculated for these different populations. For each combination of prevalence rate and population size, different number of replicates were used, ranging from 10-100 (also in increments of 10), then 200 and 500 (see Fig. S1A for a schematic representation). These simulations were all ran with the default values (i.e. successive points = 10; mean-difference = 2; Δ = 1). A example of a population with 1000 individuals, prevalence rate of 50% and 50 replicates was then taken and further parameters were tested. The number of successive points (i.e. 2, 10 and 50), mean-difference (i.e. 1, 2, 5 and 10) and Δ values (i.e. 0.5, 1 and 2) were all tested in combination with each other (Fig. S1B). One final example was taken with set parameters and settings and was run 10 times in order to evaluate the natural stochasticity of the process.

The first part of the simulations dealt with different population sizes and prevalence rates, as well as varying numbers of replicates. Default values for the other parameters revealed that, with few exceptions, all the estimated prevalence rates were within 1% of the actual prevalence rates (Table S2). The number of samples required to estimate the prevalence rate in all of our simulations was on average 32, but this value was as low as 20 for prevalence rates of 10 or 90% and as high as 40 samples for prevalence rates of 40-60%.

The second part of the simulations, where we changed the number of successive points, the value of mean-difference and the Δ value revealed more substantial variations in the results, highlighting the importance of choosing meaningful values. A higher number of successive points naturally resulted in a higher number of samples needed, as did a smaller value (i.e. 0.5) for Δ and smaller values of mean-difference (e.g. the lower the value the more samples required).

We then ran a final set of simulations 10 times using the default values for all parameters (i.e. successive points = 10; mean-difference = 2; Δ = 1; number of replicates = 50) on a population with 1000 individuals and a prevalence rate of 50%. Even though there was only one instance in which the predicted prevalence rate only varied from the actual one by more than 1%, the number of samples required to estimate it was on average 38 ± 9.5, highlighting the natural variation of the process. We therefore suggest users to run the analysis on their real data sets a minimum of five times to avoid getting a number of required samples that by chance deviates far from the mean.

### Real data: prevalence rates of coral-dwelling fauna

In order to test the accuracy of the `R` package, we ran `SAMPLE` on a data set of corals and their associated fauna, sampled from 586 host colonies at various sites along the leeward side of Curaçao (Dutch Caribbean) between the 24^*th*^ of February and the 30^*th*^ of March 2022.

Four species of stony corals (Scleractinia) and one hydrocoral (Hydrozoa) species were selected for the prediction of sampling effort needed for the accurate estimation of symbiont prevalence rate: A) the stony coral *Acropora palmata* with the crab *Domecia acanthophora* and fanworm *Spirobranchus polycerus*; B) the stony coral *Agaricia agaricites* with the crab *Opecarcinus hypostegus*; C) the stony coral *Agaricia lamarcki* with *O. hypostegus*; D) the hydrocoral *Millepora complanata* with the barnacle *Megabalanus stultus, D. acanthophora* and the blenny *Acanthemblemaria spinosa*; E) the stony coral *Orbicella faveolata* with the crab *Troglocarcinus corallicola* at 6m; and F) *Orbicella faveolata* with *T. corallicola* at 15m. This data set will enable us to detect patterns of: 1) prevalence of multiple symbionts species on a single host (A and D); 2) prevalence of the same symbiont on closely related hosts (B and C); and 3) prevalence of the same symbiont/host pair across two different depths (E and F).

Using the aforementioned coral and associated fauna data set, included in the `SAMPLE` package and in the Supplementary material (S3), we were able to estimate the minimum number of sampled corals needed to accurately estimate the prevalence rate of their symbionts. The default visualisation output of the package is a panel figure where the number of panels is equal to the number of host species from the input data set. The title of each panel corresponds to the name of the host species, and the names of the symbionts are provided in the legend at the bottom of the figure (Fig. 2). The estimated prevalence rate is provided at the right-end side of each panel, next to the tail-end of the prevalence curve, and the number of samples needed to estimate that very same prevalence rate is at the top of the panel and indicated with a dotted line, both in the same colour as the symbiont in question (see Fig. 3 for a more detailed explanation). For example, the prevalence rates of the blenny *Acanthemblemaria spinosa*, the crab *Domecia acanthophora*, and the barnacle *Megabalanus stultus* on the hydrocoral *Millepora complanata* are 21.1%, 19.2% and 23.8%, respectively. The number of sampled *M. complanata* hydrocorals needed to obtain those prevalence rates was 26, 39 and 29 (Fig. 2D), respectively, highlighting that our sampling effort (*n* = 104) was sufficient. In the case of multiple symbionts requiring the exact same number of samples, only one dotted line will be presented, but both numbers will be printed. If the number of samples is not enough to estimate the prevalence rate, then *na* is printed at the top of the graph in the colour of the respective symbiont, as can be seen for *Spirobranchus polycerus* on *Acropora palmata*. In this case `SAMPLE` indicates that our sampling effort (*n* = 35) was too low to obtain a meaningful prevalence rate for that particular symbiont (Fig. 2A). However, Fig. 2A) shows that stabilisation is close, but likely has not yet been reached because the number of successive points required surpasses the number of data points obtained.

**Fig. 3.**
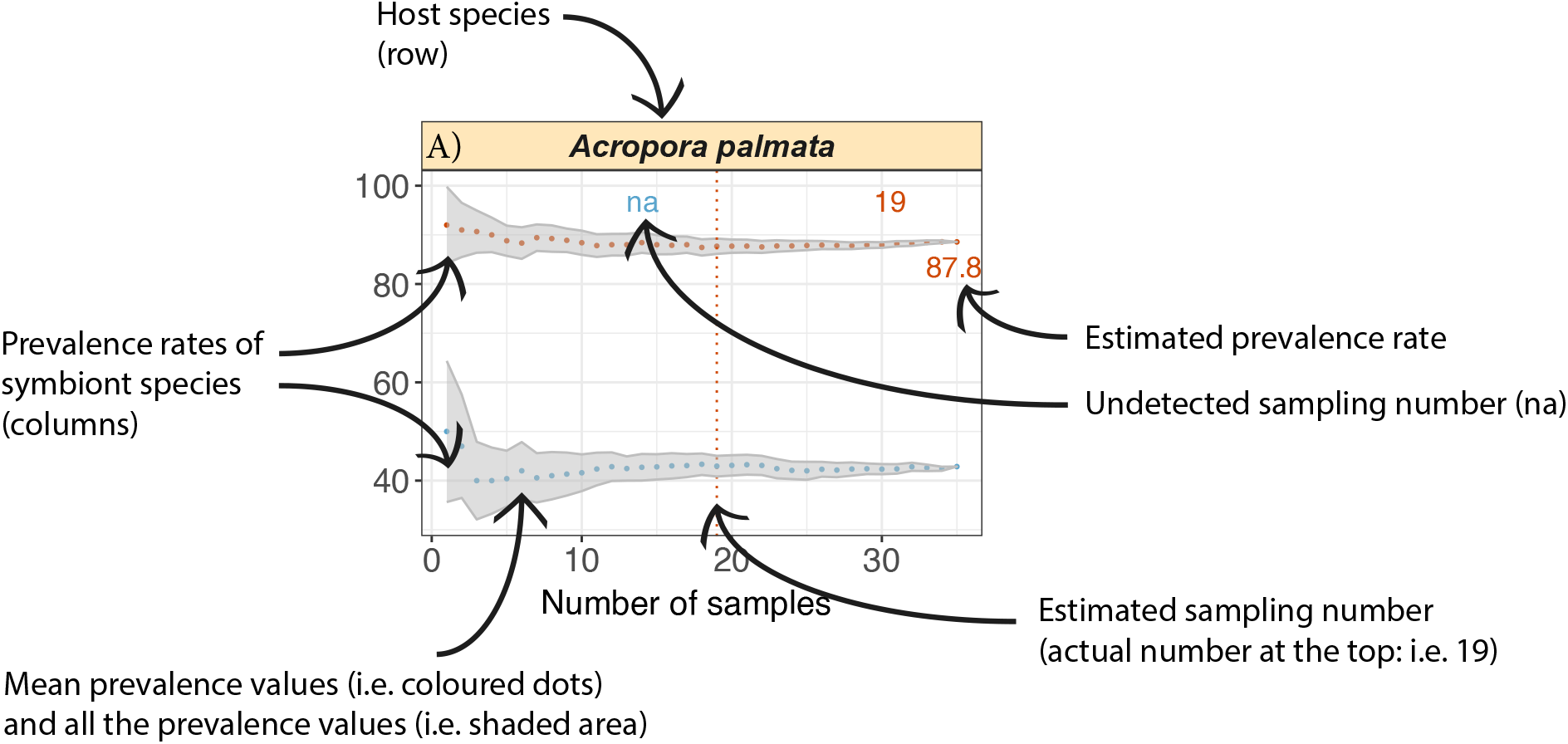
A detailed explanation of what the different elements present on the output plots represent.

## Discussion

Reliable estimations of occurrence or prevalence rates are not always easy to achieve. It has been suggested that the more samples collected, the more reliable the estimations (Gregory and Blackburn, 1991), and whilst big sample sizes tend to be preferred, this is not always necessary or possible (Underwood, 1997; Jovani and Tella, 2006). There are many factors that can contribute to small sample sizes: the sampling environment can be difficult to access (e.g. deep sea or caves, political instability in certain countries), sampling time can be limited (e.g. funding, scuba diving, boat time), or specimens are elusive and/or rare. Knowing whether the samples collected allow for a representative estimation of occurrence rates is therefore needed when sampling time and/or funds are limited.

The `SAMPLE R` package provided here aims at reducing this uncertainty and informing the user of the minimum number of samples needed to accurately estimate occurrence/prevalence rates, or otherwise informing the user that more sampling is required. `SAMPLE` can also be run during field campaigns, highlighting which taxa ideally need more data points. Even in situations where the sampling conducted is not enough to estimate a prevalence rate, and more data cannot be obtained, the visual output (see Fig. 2A as an example) allows to make a prediction as to whether a stabilisation of the prevalence rate is close or not.

This package can detect an early stabilisation of the prevalence rate of symbionts, and this is the value that is reported initially. It is possible, albeit unlikely, that another prevalence rate will be found in that same host/symbiont system that differs from the initial one if more runs are done, this is due to stochastic nature of the process. Parameters can, however, be manually changed in order to make the estimation of prevalence rate and number of samples needed more or less stringent. Moreover, small variations in prevalence rate values (i.e. less than 0.1%) are not necessarily ecologically relevant (Jovani and Tella, 2006), and the variation found in the simulation examples provided here tend to vary little (i.e. less than 1% in most cases). Ultimately it is up to the user to determine the value that is the most ecologically relevant for their particular study system and to determine the minimum sample size required to estimate it. We recommend having a careful look at the results of the simulation (see Supplementary material S2) to understand how changing certain parameters may affect the estimated prevalence rates and the number of samples required and then, if necessary, adjust the parameters to fit specific needs.

One of the strengths of this `R` package is also one of its weaknesses: being able to set so many parameters allows for users to choose the properties that suit them best, leading to robust results. On the other hand, it also means that if users do not know their study species or system well, they might end up choosing parameters that are not the most suitable. The default values should, however, be suitable for most study systems. The nature of this bootstrapping approach also comes with some intrinsic variation, so there will be some stochasticity in the results. As mentioned before, a way of counteracting this is by repeatedly running the analysis (we suggest a minimum of five runs) so the results obtained can converge, but there will always be some variation. Lastly, some prevalence rates can be harder to estimate, particularly those that are very high or very low (Gregory and Blackburn, 1991; Jovani and Tella, 2006), as highlighted by the results of our simulation, so when possible, users should be less conservative with their sampling effort to account for this.

It is important to note that prevalence rates of symbionts across their hosts, even when looking at the same host/symbiont system, can change across time and space (Shykoff and Kaltz, 1998; Penczykowski et al., 2016; Starkloff and Galen, 2023). In the example provided above, we looked at the same species of symbiont, the crab *T. corallicola*, on the same coral host, *O. faveolata*, across two different depths (6 and 15 m). `SAMPLE` detected different prevalence rates, and slightly different number of samples needed (*n* = 20 and 25, respectively) for the estimation of prevalence rates (Fig. 2E-F). We also estimated the prevalence rates of the same symbiont, the crab *O. hypostegus*, on two closely related coral host species, *A. agaricites* and *A. lamarcki*, across the same depth range (i.e. from 6 to 15 m) and the results varied considerably (Fig. 2B-C). These examples highlight that ecological factors impact symbiont prevalence rates across different depths and hosts (van Tienderen and van der Meij, 2016). Ultimately it is up to the user to choose when and how to use this package to best suit their needs.

Finally, it should also be mentioned that this package is not aimed at determining overall prevalence rates for host/symbiont systems, or overall occurrence rates of species. It is rather aimed at estimating if the sampling effort is enough to estimate a stable rate for that particular group of taxa in that particular point in time.

## Acknowledgements

The data collection in the field was a joint effort conducted by the first author, Natascha Borgstein, Tao Xu and Yun Scholten (all University of Groningen). Without their help there would be no real data set for this study. We thank Pedro Santos Neves and Raphaël Scherrer, also from the University of Groningen, for their help in compiling the original code into the `SAMPLE R` package. We further thank the Center for Information Technology of the University of Groningen for their support and for providing access to the Hábrók high performance computing cluster. Funding for fieldwork in Curaçao was financed by Academy Ecology Fund Grant from the Royal Netherlands Academy of Arts and Sciences (No. KNAWWF/705/ECO202223), TREUB-maatschappij (Society for the Advancement of Research in the Tropics), FONA Foundation for Research on Nature Conservation, and Stichting Fonds Dr. Christine Buisman. A final thank you to the CARMABI research station on Curaçao for helping us with logistical support.

## Declarations

### Competing Interests

The authors have no competing interests to declare that are relevant to the content of this article.

#### Data Availability

All the data and scripts used in this study are available in the following online repository: https://github.com/yacinebenchehida/SAMPLE/

## Supplementary Material

Supplementary material not included in this section can be found in the following online repository: https://github.com/yacinebenchehida/SAMPLE/

**Fig. S1.**
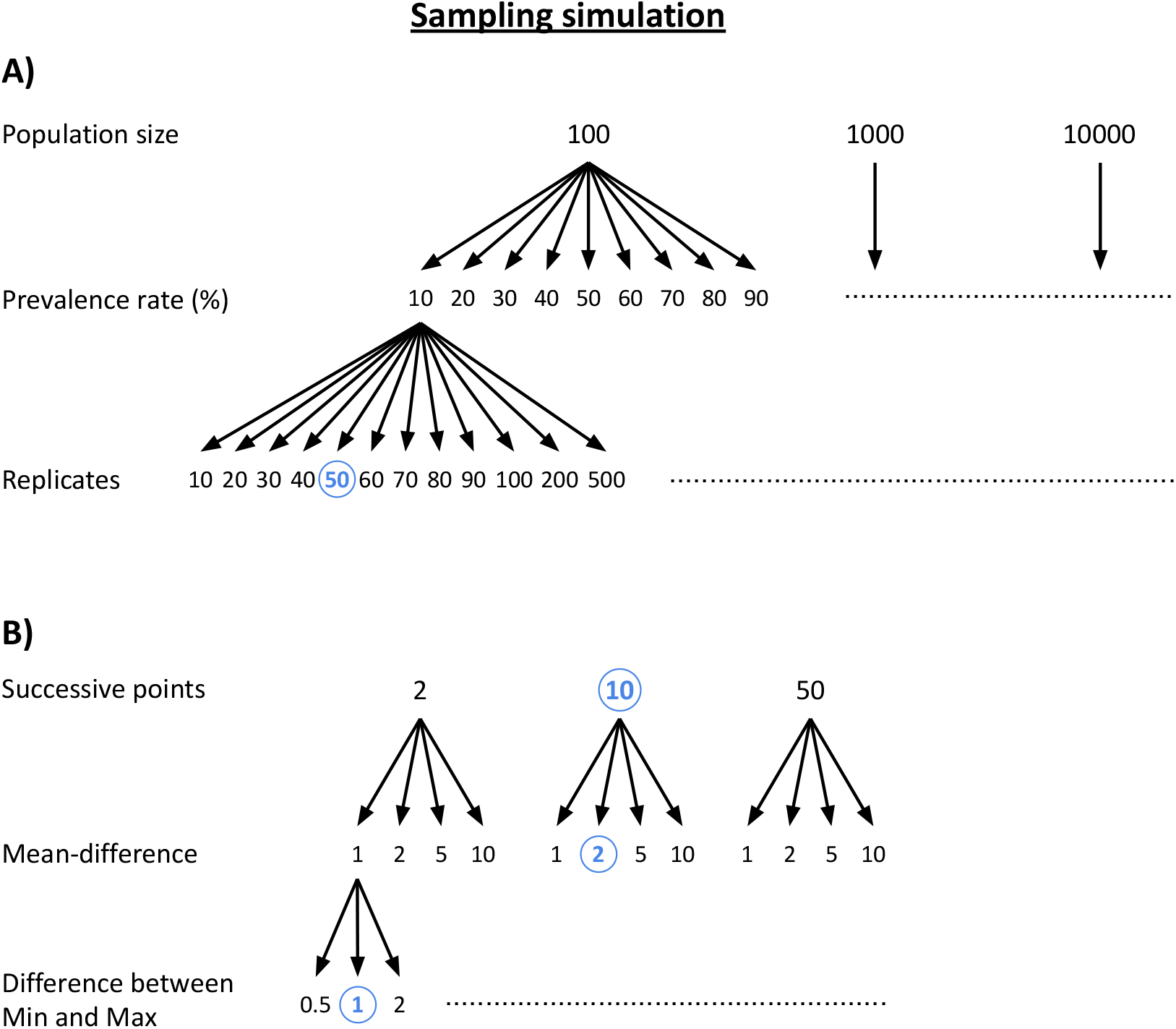
Schematic representation of the simulation sampling, where: different sized populations were sampled with different prevalence rates and different number of replicates **(A)**; and one example of a population with 1000 individuals, a prevalence rate of 50% and 50 replicates was then taken and ran with different settings of successive points, mean-difference and Δ **(B)**. A final example was chosen and ran 10 times with the exact same settings and using the default values highlighted in blue in order to evaluate the natural variation in the process.

