## Supplementary figures and images for "SAMPLE: an R package to estimate sampling effort for species’ occurrence rates"

### Supplementary material S1

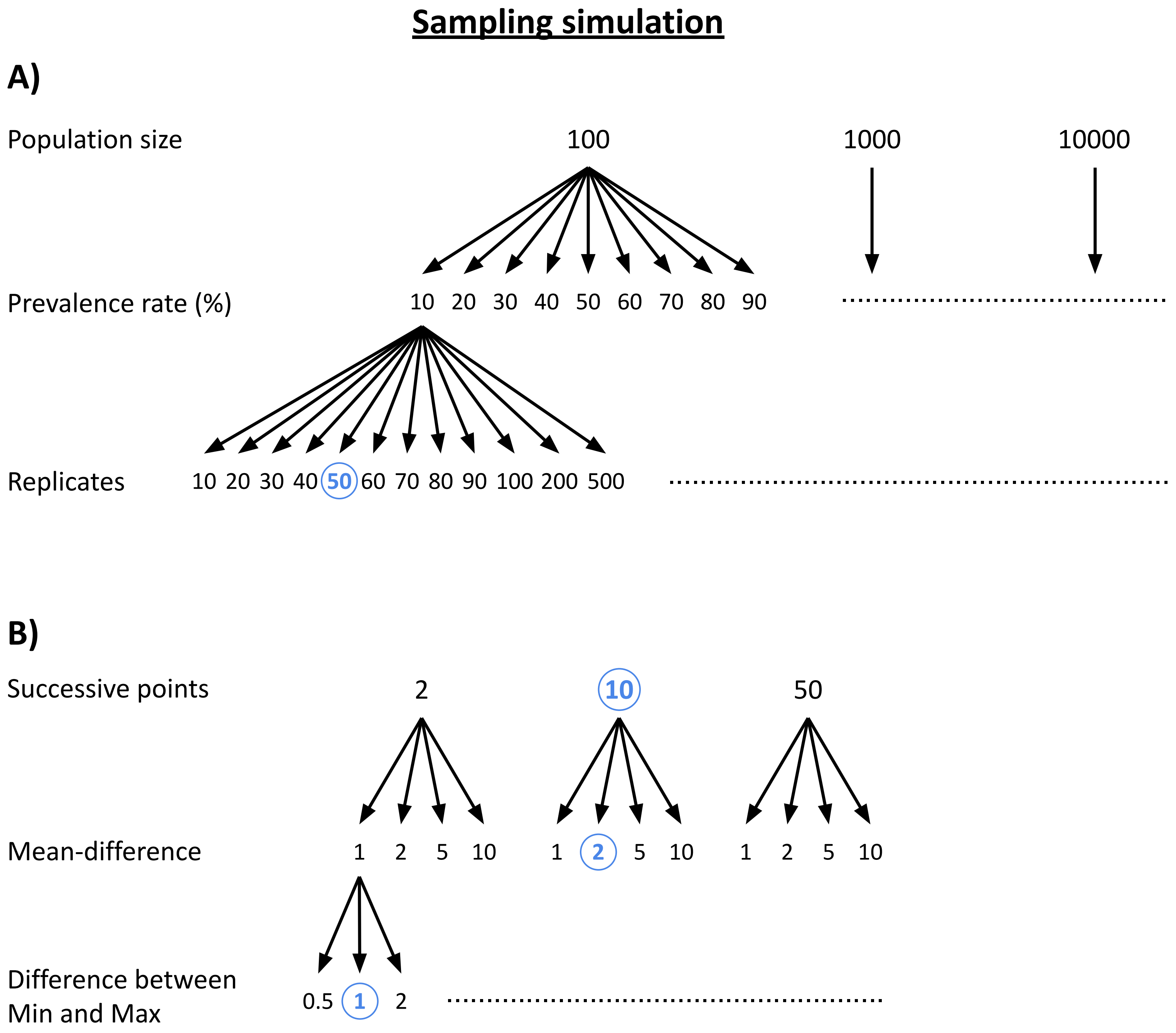
